# Chronic Developmental Lead Exposure increases μ-Opiate Receptor Levels in the Adolescent Rat Brain

**DOI:** 10.1101/2020.04.18.048264

**Authors:** Damaris Albores-Garcia, Jennifer L McGlothan, Zoran Bursac, Tomás R. Guilarte

## Abstract

Opioid use and abuse has reached epidemic proportion in the United States resulting in a significant numbers of deaths due to overdose. While environmental factors are implicated in opioid addiction, less is known about the role of exposure to environmental pollutants on the brain opioid system. Human and preclinical studies have suggested an association between childhood lead (Pb^2+^) intoxication and proclivity to substance abuse and delinquent behavior. Opioid receptors are involved in the biological effects of opioids and other drugs of abuse. In this study, we examine the effect of chronic developmental Pb^2+^ exposure on μ-opioid receptor (MOR) levels in the rat brain using [^3^H]-D-Ala2-MePhe4-Gly-ol5 enkephalin ([^3^H]-DAMGO) quantitative receptor autoradiography.

Our results indicate that chronic developmental Pb^2+^ exposure increases the levels of [^3^H]-DAMGO specific binding to MOR in several limbic regions of the brain in male and female rats during the pre-adolescence (PN14) and early-adolescence (PN28) period. These changes were less pronounced in late-adolescence (PN50) and adult (PN120) animals. Our findings are important because the pre-adolescence and early adolescence period is a time in which there is higher engagement in reward and drug seeking behaviors in humans.

In summary, we show that chronic exposure to Pb^2+^ an ubiquitous and well-known environmental contaminant and neurotoxicant, alters MOR levels in brain regions associated with addiction circuits in the adolescent period with important implications to opioid drug use and abuse.

## 1. Introduction

Childhood lead (Pb^2+^) intoxication is a public health problem of major proportion throughout the world (Dooyema et al., 2012; Li et al., 2014; Olympio et al., 2017; Raymond and Brown, 2016; Zhang et al., 2020). In the United States, public health policy implemented in the late 1970s to remove Pb^2+^ from paint and gasoline was effective in markedly reducing the overall levels of Pb^2+^ in blood of the general population and in particular in children (Lanphear et al., 2003; Pirkle et al., 1994; Pirkle et al., 1998). However, there is still a significant number of children with blood Pb^2+^ levels that are detrimental to their health (Christensen et al., 2019; Han et al., 2018; Neuwirth, 2018; Tirima et al., 2018). Recent events indicate that a significant number of municipalities suffer from Pb^2+^ contamination in drinking water in homes and schools throughout the United States (Hanna-Attisha et al., 2016). Childhood Pb^2+^ intoxication is most likely to occur in low-income, inner city communities where there is a higher number of Pb^2+^ contaminated homes and decaying infrastructure with a disproportionate burden in disadvantage African American and Hispanic communities (Cassidy-Bushrow et al., 2017; Rothweiler et al., 2007; White et al., 2016; Yeter et al., 2020).

Childhood Pb^2+^ intoxication is best known for producing deficits in learning and poor school performance (Baghurst et al., 1992; Chen et al., 2005; Dietrich et al., 1993; Plusquellec et al., 2007). Furthermore, there is evidence that childhood Pb^2+^ intoxication is associated with delinquent and maladaptive behaviors as well as increased use of drugs of abuse later in life (Desrochers-Couture et al., 2019; Fishbein et al., 2008; Needleman et al., 1996; Wright et al., 2008). In the last decades, preclinical and population-based studies have provided evidence of an association between chronic developmental Pb^2+^ exposure and mental disorders such as major depression, mood disorders, and schizophrenia (Abazyan et al., 2014; Guilarte et al., 2012; Kponee-Shovein et al., 2020; Stansfield et al., 2015). Preclinical studies indicate that developmental Pb^2+^ exposure may be associated with propensity to opiate addiction (Rocha et al., 2004). A study in inner city woman that abuse heroin showed a nearly doubling in tibial Pb^2+^ concentrations using x-ray fluorescence (Fishbein et al., 2008). Furthermore, studies originating in Iran indicate that opium users have higher blood Pb^2+^ levels than non-users, although this may be due to the potential Pb^2+^ contamination of the heroin preparation (Amiri and Amini, 2012; Salehi et al., 2009; Soltaninejad and Shadnia, 2018).

In the last decade, opioid use disorder has reached epidemic proportions in the United States with a highly significant increase in overdose deaths from opioids, synthetic opioids, and heroin (Wilson et al., 2020). The Center of Disease Control and Prevention estimates nearly 68,000 drug overdose deaths in 2018 (Wilson et al., 2020). Furthermore, inner city low-income populations are more likely to misuse opioids and have opioid use disorder than the general population (Altekruse et al., 2020). This is the same group that is most likely to be exposed to environmental toxins such as Pb^2+^.

In the present study we examined the effect of chronic developmental Pb^2+^ exposure on the brain opioid system in an effort to understand a possible relationship between childhood Pb^2+^ intoxication and the current opioid epidemic. To this aim, we examine the effect of developmental Pb^2+^ exposure on the levels of mu (μ) opiate receptors (MOR) in the rat brain using quantitative receptor autoradiography. In the brain, MOR is involved in reward-seeking behaviors and play an important role in opioid use disorders (Darcq and Kieffer, 2018; Gerrits et al., 2003) and positive reinforcement of opioids as well as non-opioid drugs of abuse (Contet et al., 2004). Our results indicate that developmental Pb^2+^ exposure results in increased levels of MOR in regions of the brain that play an important role in drug addiction and these changes are specific to the adolescent period with important implications to opioid use and abuse.

## 2. Materials and Methods

### 2.1 Animal Husbandry

Adult female Long-Evans rats (225-250g; Charles River Inc., Wilmington, MA, USA) were fed a diet containing either 0 or 1500 parts per million (ppm) Pb^2+^-acetate (RMH 1000; Dyets, Bethlehem, PA, USA) starting 10 days prior to breeding to non-exposed males and throughout gestation and lactation. One day after birth, litters were culled to 10 pups and weaned twenty-one days after birth (PN-21). After weaning, animals were fed the same diet as their respective dam in order to maintain a continuous Pb^2+^ exposure throughout life. At fourteen (PN14: pre-adolescence), twenty-eight (PN28: early adolescence), fifty (PN50: late adolescence), or one hundred and twenty days after birth (PN120: adult), male and female offspring were euthanized, blood collected, and the brain harvested and kept at −80°C until used. The definition of the different periods of development used in this manuscript were adapted from Spear (Spear, 2000). For statistical purposes, a single data point consisted of one male or female pup per litter, making the litter the statistical unit. All animal studies were approved by the Florida International University Animal Care and Use Committee and have been carried out in agreement with the Guide for Care and Use of Laboratory Animals as stated by the U.S. National Institutes of Health.

### 2.2 Blood Pb^2+^ levels

Rats were euthanized at the appropriate age using CO_2_ and the blood samples were collected transcardially and placed into 1 cc vacutainer EDTA tubes. Fifty μL of the blood sample was added to the LeadCare plus tubes and blood Pb^2+^ levels were measured 24 to 72 hours after, using a Magellan LeadCare plus analyzer following manufacturer’s instructions (ESA Laboratories, Chelmsford, MA, USA).

### 2.3 Tissue Collection and Quantitative Autoradiography

At the different ages, rat brains were harvested immediately after decapitation, rinsed with saline and frozen in dry ice and stored at −80°C until used. Fresh-frozen brains were sectioned at 20-micron thickness in the coronal plane with a freezing cryostat (Leica SM2000R; Leica Microsystems, Wetzlar, Germany) and thaw-mounted on poly-L-lysine-coated slides. Slides were stored at −80°C until used.

For μ opioid receptor autoradiography, slides with brain slices were thawed and pre-incubated in 50 mM Tris buffer + 0.9 % NaCl (pH 7.4) at room temperature for 30 minutes. For total binding, slides were incubated in Tris buffer with 3.9-4.1 nM [^3^H]-D-Ala^2^-MePhe^4^-Gly-ol^5^ enkephalin ([^3^H]-DAMGO) in ice-cold Tris-HCl buffer (pH 7.4) for 60 minutes at room temperature. Non-specific binding was assessed in adjacent brain slices incubated in the same solution and conditions as for total binding but with the addition of 1 μM of the opioid antagonist naloxone. Following incubation with [^3^H]-DAMGO, slides were rinsed three times in buffer at 4°C and then dipped once in dH_2_O at 4°C. After drying overnight, slides were apposed to Kodak Biomax MR film, MR-1, for 16 weeks. [^3^H]-Microscales (Amersham, Arlington Heights, IL, USA) were included with each film to allow for quantitative analysis of images. All brain slices (total and non-specific) from a particular age were placed in one film so that brain slices from PN14, PN28, PN50, and PN120 were all in different films. Following the 16-week period, films were developed, and images were captured and analyzed using MCID Imaging software (MCID, InterFocus Imaging, Cambridgeshire, UK). A rat brain atlas [Paxinos and Watson, 1998] was used to define brain regions. In any one brain region, specific binding was determined by subtracting the non-specific binding from the total binding of [^3^H]-DAMGO to MOR. Non-specific binding was essentially zero or less than 1% of total binding. Therefore, with [^3^H]-DAMGO total binding is essentially specific binding.

### 2.4 Statistical Analysis

Statistical analyses were performed with SAS/STATv14.2 (SAS Institute Inc., Cary, NC) and GraphPad Prism 8. Outcome variable was examined for outliers and normality using a Kolmogorov-Smirnov test. Summary statistics including means, standard deviations, frequencies, and proportions were generated for main outcome MOR and independent experimental conditions, treatment, sex, age and brain region, respectively. Summary statistics for MOR were further generated for all levels of independent experimental conditions. Univariate comparisons for MOR across experimental conditions consisted of t-test and ANOVA followed by Bonferroni adjusted multiple comparisons, depending on the number of categories in each condition, respectively. Sensitivity analysis was performed using the non-parametric equivalent tests, Wilcoxon Mann-Whitney U and Kruskal-Wallis, respectively. To test all experimental conditions in one general linear model we applied linear regression starting with all four conditions. We tested two-way treatment interactions with age, sex and brain region as well as a three-way interaction of treatment with age and sex. Instances where interactions were significant warranted further stratified investigation of simplified models. All associations were considered significant at the alpha level of 0.05. All values are expressed as mean ± S.E.M.

## 3. Results

### 3.1 Blood Pb^2+^ concentrations and body weight

The Pb^2+^-exposure paradigm used in this study did not change the body weight gain of animals at any of the ages evaluated from early-adolescence to adults; that is at PN14, PN28, PN50 and PN120 (Table 1). Blood lead levels (BLL) were significantly higher in all the Pb^2+^-exposed male and female rats at the four ages examined (Table 1). N values expressed the number of litters evaluated.

**Table 1:** Body weight and blood lead levels (μg/dL) of control and Pb^2+^-exposed male and female rats as a function of age. Data are expressed as the mean ± S.E.M.

| | Body Weight (g)<br>Mean ± SEM | | | | | | | |
| --- | --- | --- | --- | --- | --- | --- | --- | --- |
|  | PN14 |  | PN28 |  | PN50 |  | PN120 |  |
|  | Males | Females | Males | Females | Males | Females | Males | Females |
| **Control** | 34.7 ± 1<br>n=10 | 35.7 ± 1.4<br>n=4 | 120.3 ± 6.2<br>n= 10 | 109.9 ± 3.6<br>n=13 | 312 ± 10.8<br>n=10 | 224 ± 3.9<br>n=15 | 681.6 ± 16<br>n=14 | 401 ± 12.2<br>n= 13 |
| **Pb<sup>2+</sup>** | 38.2 ± 1.6<br>n=7 | 43.7 ± 3.7<br>n=4 | 112.2 ± 3.3<br>n= 13 | 109.5 ± 2.8<br>n=14 | 315 ± 10.64<br>n=9 | 214.1 ± 3.4<br>n=13 | 659 ± 8.8<br>n=20 | 400 ± 7.9<br>n= 15 |
| **P-value** | **0.0873** | **0.0846** | **0.2293** | **0.9337** | **0.8479** | **0.0736** | **0.1951** | **0.9288** |

| Blood Lead Levels ($\mu\text{g/dL}$)<br>Mean ± SEM | | | | | | | | |
| --- | --- | --- | --- | --- | --- | --- | --- | --- |
|  | PN14 |  | PN28 |  | PN50 |  | PN120 |  |
|  | Males | Females | Males | Females | Males | Females | Males | Females |
| **Control** | $\leq 1.9$<br>n=6 | $\leq 1.9$<br>n=3 | $\leq 1.9$<br>n=18 | $\leq 1.9$<br>n=16 | $\leq 1.9$<br>n=17 | $\leq 1.9$<br>n=15 | $\leq 1.9$<br>n=17 | $\leq 1.9$<br>n=15 |
| **Pb<sup>2+</sup>** | $36.1 \pm 2.9$<br>n=8 | $37 \pm 3.8$<br>n=4 | $21.1 \pm 0.9$<br>n=18 | $20.9 \pm 1.1$<br>n=16 | $20.2 \pm 1.1$<br>n=17 | $22.1 \pm 1.25$<br>n=20 | $19.6 \pm 1.3$<br>n=17 | $24.3 \pm 2.2$<br>n=14 |
| **P-value** | $\leq 0.0001$ | $\leq 0.0001$ | $\leq 0.0001$ | $\leq 0.0001$ | $\leq 0.0001$ | $\leq 0.0001$ | $\leq 0.0001$ | $\leq 0.0001$ |

### 3.2 Ontogeny of μ-opioid receptor levels in the normal rat brain

#### 3.2.1 Age effect

In control animals, analysis of MOR levels changes at different ages show that the peak of MOR expression in the rat brain, regardless of sex and brain region, is in late adolescent rats, i.e. PN50 animals (Figure 1A). There are no differences in MOR levels between pre-adolescence rats (PN14: 53.3 ± 2.7 fmol/mg tissue) and early-adolescent rats (PN28: 55.2 ± 2.5 fmol/mg tissue). There is a highly significant increase in MOR levels in late adolescent rats (PN50: 88.6 ± 2.5 fmol/mg of tissue; p<0.0001) compared to pre- and early-adolescent rat, and there is a statistically significant decrease of MOR levels in adults (PN120: 72.9 ± 2.7 fmol/mg of tissue; p=0.0001) compared to late adolescent rats (PN50). Finally, in controls rats MOR levels did not show a gender difference at any of the ages when all brain regions were included (Figure 1B).

**Figure 1:**
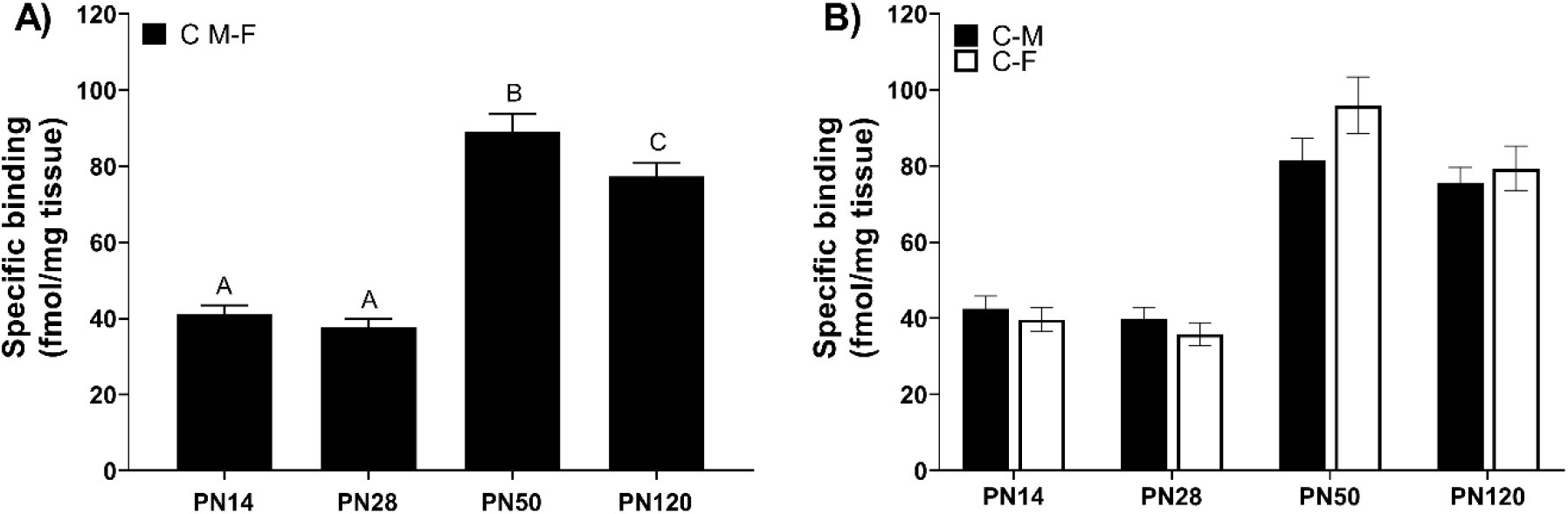
[^3^H]-DAMGO specific binding to μ-opioid receptor in the brain. A) [^3^H]-DAMGO specific binding to rat brain from combined male and female control rats at different ages. In panel A, bars with different letters are significantly different at (p<0.0001). B) [^3^H]-DAMGO specific binding in male vs female rats at the different ages including all brain regions. Values are mean ± S.E.M. n= 6-8 animals/experimental group.

#### 3.2.2 Effect of age and sex

Figure 2 shows representative images of the [^3^H]-DAMGO specific binding in control male and female rats at all ages measured in the study. When evaluated by brain region (Table 2), PN28 control males showed an increase in MOR levels compared to their female counterparts in NACc (p=0.025), and hypothalamus (p=0.0005); while females showed increase MOR levels in medial thalamus at PN28 and PN50 (p =0.0005, p=0.048, respectively).

**Table 2:** [^3^H]-DAMGO specific binding to μ-opioid receptor in different brain regions in control male and female rats as a function of age. (Hypothalamus, Thalamus, Basolateral amygdala -*BLA*-, Striatum, Lateral post-thalamic nuclei -*LPTN*-, stria medullaris of the thalamus -*SM*-, Striatum, Nucleus Accumbens). Statistical differences are indicated. Values are mean ± S.E.M. n= 6-8 animals/experimental group.

|  | PN14 | PN28 | PN50 | PN120 |
| --- | --- | --- | --- | --- |
|  | Control-M n=5-7<br>Control-F n=5-6 | Control-M n=7<br>Control-F n=7-8 | Control-M n=7<br>Control-F n=7-8 | Control-M n=7<br>Control-F n=6 |
| **MedialThalamus** |  |  |  |  |
| Control-Males | 40.9 ± 8.8 | 11.85 ± 5.6 | 131.9 ± 7.2 | 108.9 ± 7.8 |
| Control-Females | 41.8 ± 12.7 | 48.4 ± 5.9 | 154.2 ± 7.16 | 93.4 ± 4.00 |
|  | ns | **p=0.0005** | **p=0.048** | ns |
| **Hypothalamus** |  |  |  |  |
| Control-Males | 42.6 ± 6.0 | 62.7 ± 8.0 | 47.4 ± 6.2 | 30.07 ± 3.94 |
| Control-Females | 36.7 ± 6.5 | 5.7 ± 2.2 | 36.9 ± 4.7 | 25.9 ± 1.7 |
|  | ns | **p<0.0001** | ns | ns |
| **SM**<br>Control-Males<br>Control-Females | 52.4 ± 15.0<br>60.8 ± 14.08<br>ns | 49.3 ± 7.1<br>51.3 ± 12.7<br>ns | 133.5 ± 24.7<br>194.0 ± 23.1<br>ns | 85.1 ± 13.9<br>127.1 ± 33.5<br>ns |
| **LPTN**<br>Control-Males<br>Control-Females | 30.8 ± 6.7<br>31.6 ± 11.4<br>ns | 39.0 ± 6.3<br>35.2 ± 7.3<br>ns | 94.8 ± 9.8<br>120.9 ± 8.8<br>ns | 66.9 ± 7.2<br>64.1 ± 6.7<br>ns |
| **BLA**<br>Control-Males<br>Control-Females | 46.9 ± 5.0<br>43.9 ± 9.7<br>ns | 20.6 ± 7.1<br>20.84 ± 7.36<br>ns | 77.1 ± 10.5<br>78.8 ± 6.2<br>ns | 68.0 ± 9.7<br>73.3 ± 8.8<br>ns |
| **Striatum**<br>Control-Males<br>Control-Females | 36.9 ± 7.2<br>40.0 ± 5.0<br>ns | 43.7 ± 7.4<br>47.5 ± 3.7<br>ns | 61.1 ± 5.0<br>67.8 ± 6.4<br>ns | 88.9 ± 6.5<br>86.3 ± 7.5<br>ns |
| **NACc**<br>Control-Males<br>Control-Females | 33.5 ± 5.7<br>34.9 ± 4.6<br>ns | 50.9 ± 4.1<br>36.3 ± 3.8<br>**p=0.025** | 54.9 ± 4.9<br>61.3 ± 9.7<br>ns | 75.4 ± 5.9<br>78.2 ± 5.8<br>ns |
| **NACs**<br>Control-Males<br>Control-Females | 39.9 ± 1.2<br>32.6 ± 4.8<br>ns | 46.0 ± 3.6<br>44.3 ± 2.2<br>ns | 51.9 ± 5.2<br>67.1 ± 8.4<br>ns | 81.2 ± 10.3<br>87.0 ± 6.9<br>ns |

**Figure 2:**
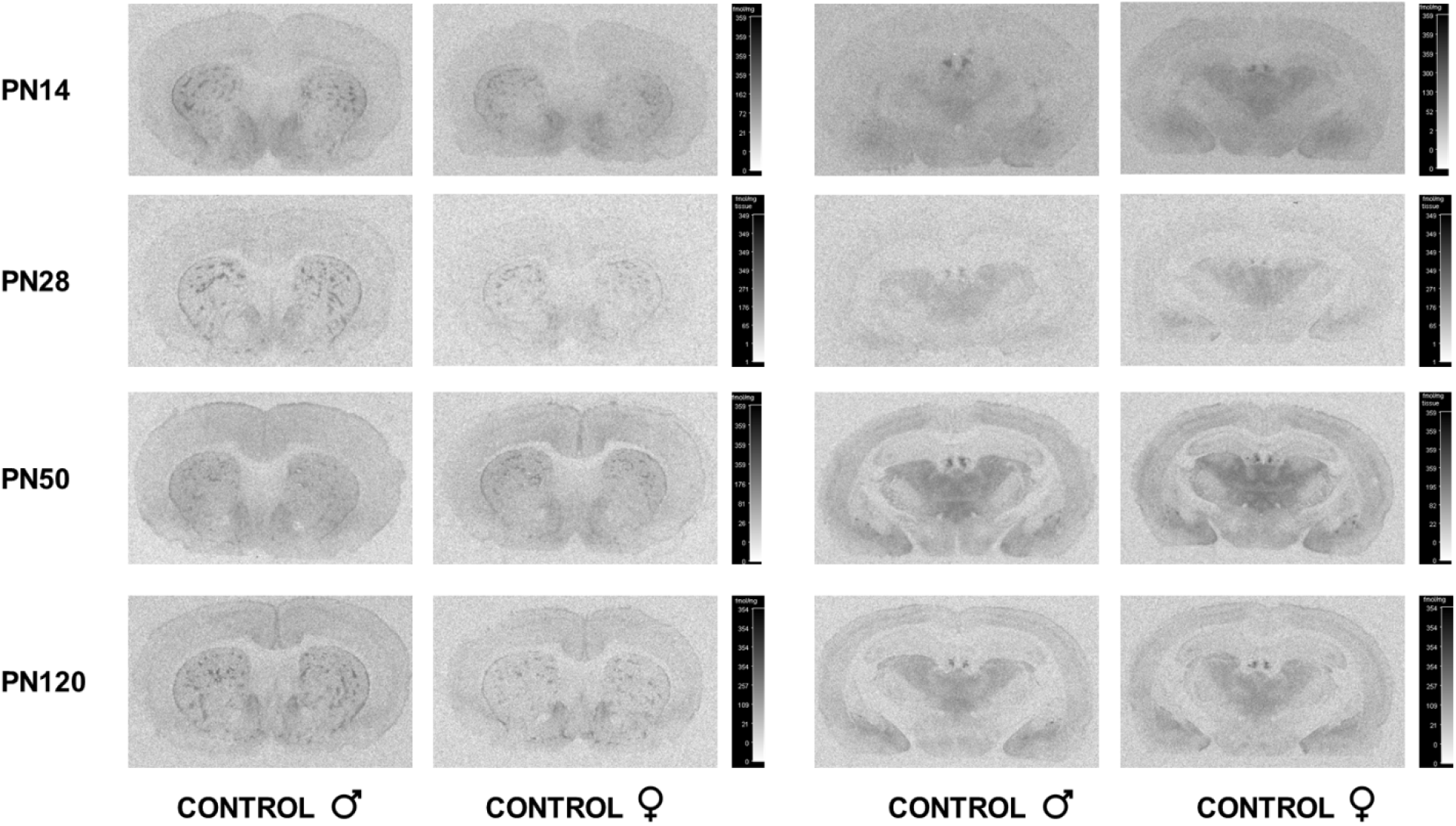
[^3^H]-DAMGO autoradiography. Representative images of [^3^H]-DAMGO specific binding to μ-opioid receptors in different brain regions of male and female control rats as a function of age.

### 3.3 Chronic developmental Pb^2+^ exposure alters μ-opioid receptor levels in the rat brain in an age-dependent fashion

We measured [^3^H]-DAMGO specific binding to MOR in several limbic regions from control and Pb^2+^-exposed rat brain using a life course approach. Overall, chronic developmental Pb^2+^-exposure had a statistically significant increase (p <0.0001) on MOR levels in the rat brain assessed by [^3^H]-DAMGO specific binding regardless of age, sex, and brain region analyzed (Figure 3A). When analyzed by age and sex (regardless of brain region), we found an age-dependent effect of Pb^2+^-exposure, showing overall higher levels of [^3^H]-DAMGO specific binding in Pb^2+^-exposed male and female rat brain at earlier developmental stages (PN14-juvenile and PN28-early adolescence) with no effect at PN50-late adolescence and a significant decrease in male Pb^2+^ exposed animals at PN120-adults (Figure 3B and 3C represents data from males and females, respectively).

**Figure 3:**
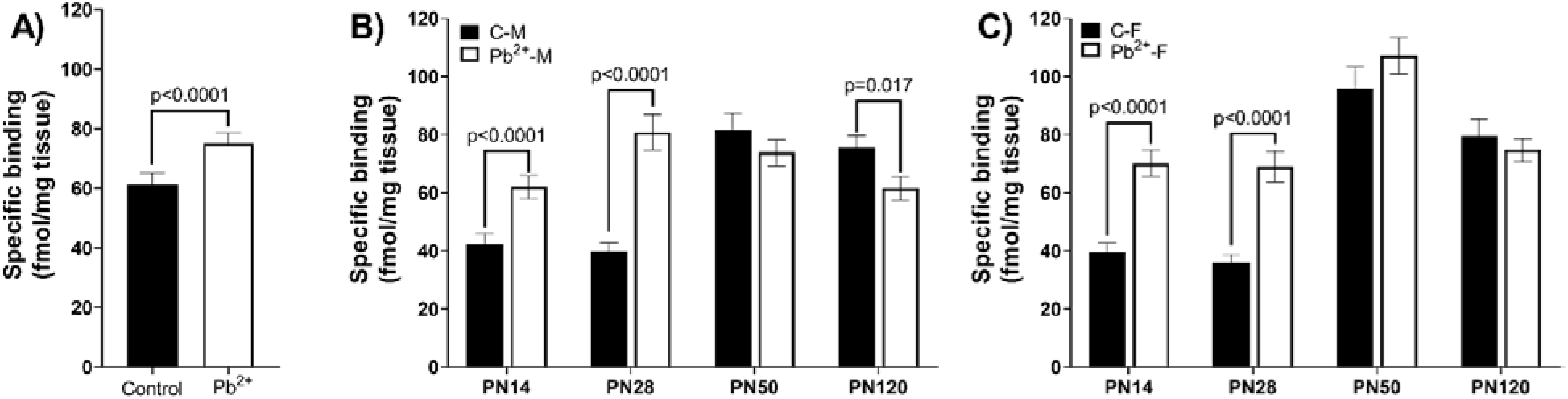
[^3^H]-DAMGO specific binding to μ-opioid receptor in the rat brain of control and Pb^2+^-exposed rats. A) Specific binding of [^3^H]-DAMGO to μ-opioid receptor in control vs Pb^2+^-exposed rats (including sex, ages and brain regions evaluated). B) [^3^H]-DAMGO specific binding to μ-opioid receptor in the brain of male control and Pb^2+^-exposed rats at PN14, PN28, PN50 and PN120. C) [^3^H]-DAMGO specific binding to μ-opioid receptor in the brain of female control and Pb^2+^-exposed rats at all ages. Values are mean ± S.E.M. n= 6-8 animals/experimental group.

We then examined the effect of chronic Pb^2+^ exposure [^3^H]-DAMGO to MOR based on brain region and sex. Figure 4 provides representative autoradiograms of [^3^H]-DAMGO to MOR of the different brain regions and ages of controls and Pb^2+^-exposed male rats. <u>Pre-adolescence</u>: At PN14, Pb^2+^-exposed male rats had higher MOR levels in medial-thalamus (p=0.018), lateral post-thalamic nuclei (LPTN) (p= 0.007), and nucleus accumbens shell (NACs) (p=0.015). While in PN14 females, Pb^2+^-exposed rats had increased MOR levels in medial-thalamus (p=0.006), hypothalamus (p=0.0125), LPTN (p=0.001), basolateral amygdala (BLA) (p=0.047), and stria medullaris of the thalamus (SM) (p=0.006) (Figure 5). <u>Early adolescence</u>: In PN28 male rats, Pb^2+^-exposed rats had an increase in MOR levels in the hypothalamus (p=0.0015), LPTN (p=0.0032), BLA (p=0.0024), STR (p=0.0075), NACc (p=0.0395), nucleus accumbens shell (NACs) (p=0.0137), and SM (p=0.0026). For PN28 females, Pb^2+^-exposure increase the MOR levels in the medial-thalamus (p=0.0023), hypothalamus (p=0.0344), LPTN (p=0.0064), BLA (p=0.042), STR (p=0.0445), NACc (p=0.0052) and NACs (p=0.0295) (Figure 5). <u>Late adolescent</u>: At PN50, Pb^2+^-exposure did not cause any changes in the MOR levels in any brain region analyzed from male rats compared to their control counterparts (Figure 5). In PN50 females, Pb^2+^-exposed females had higher levels of MOR in STR (p=0.042), whereas no other changes were observed. <u>Adult</u>: In adult rats (PN120), male Pb^2+^-exposed rats showed a decrease in MOR levels in the medial-thalamus compared to controls (p=0.0189). No changes in MOR levels were observed in female rats.

**Figure 4:**
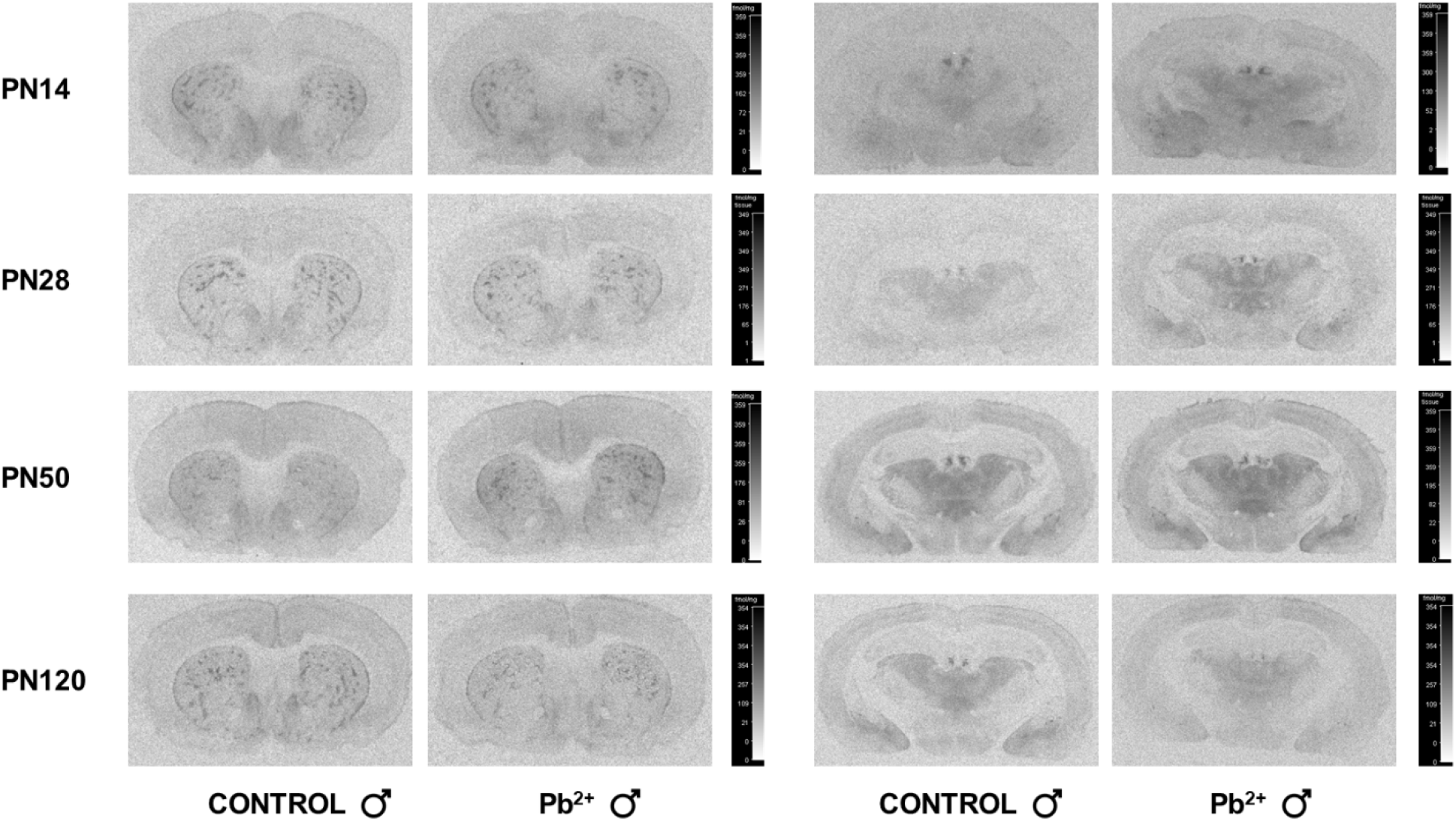
Representative images of [^3^H]-DAMGO autoradiography showing the distribution and levels of μ-opioid receptors in different brain regions in control and Pb^2+^-exposed male rats at PN14, PN28, PN50 and PN120.

**Figure 5:**
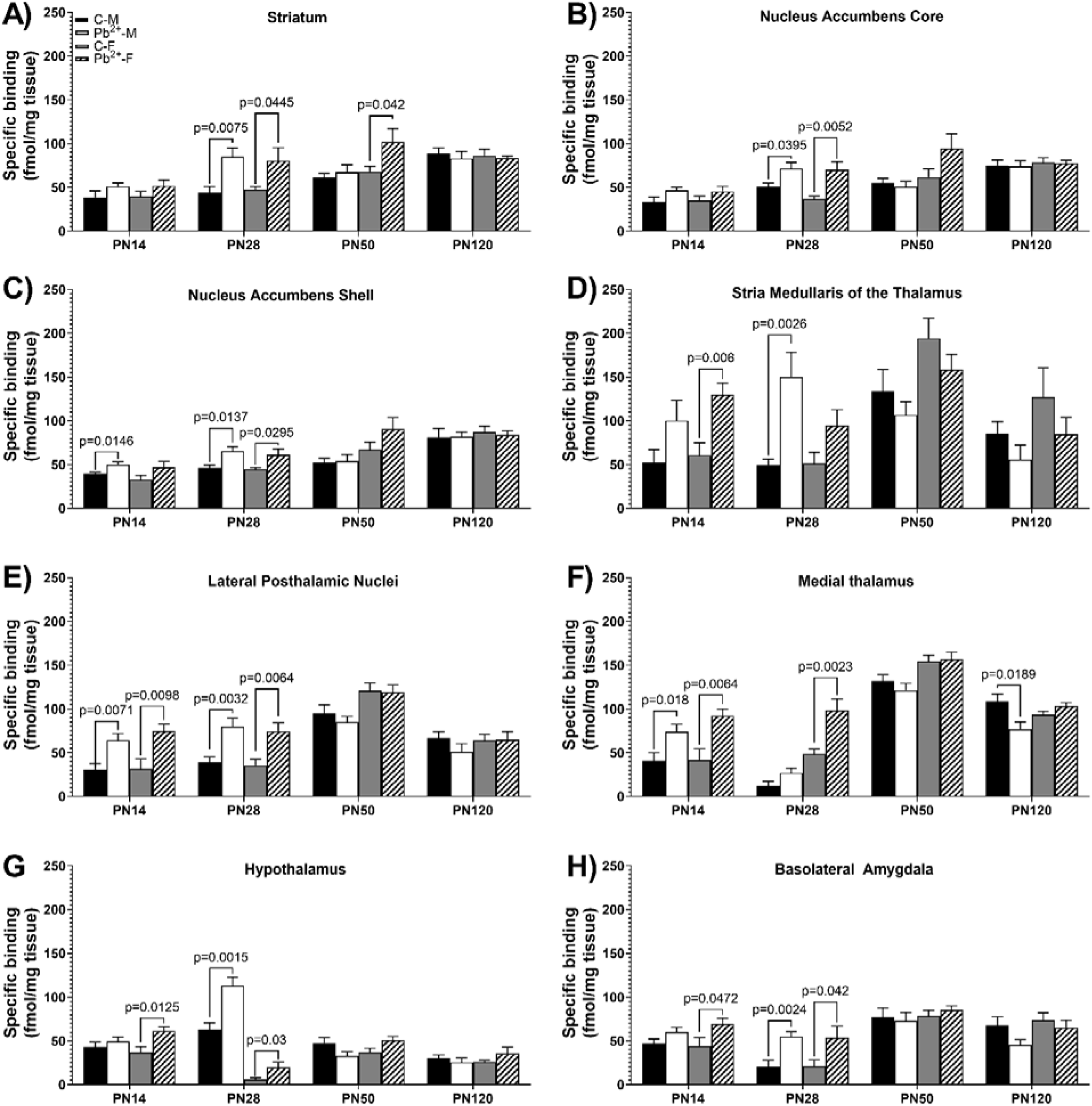
[^3^H]-DAMGO specific binding to μ-opioid receptor in different brain regions in control and Pb^2+^-exposed male and female rats as a function of age. A) striatum, B) nucleus accumbens core, C) nucleus accumbens shell, D) stria medullaris of the thalamus, E) lateral posthalamic nuclei, F) medial thalamus, G) hypothalamus, and H) basolateral amygdala. Statistical differences compared to their correspondent control group are indicated. Values are mean ± S.E.M. n= 6-8 animals/experimental group.

## 4. Discussion

The main finding of the present study is that chronic developmental Pb^2+^ exposure increased MOR levels in several limbic regions of the brain in pre-adolescence and early-adolescence male and female rats. As animals aged beyond the juvenile period (PN28-42), the effects of Pb^2+^ on MOR levels were more subtle, showing no effect on late adolescence (PN50) males and an increase in only one brain region in Pb^2+^-exposed late adolescence females (Figure 5). Further, there were no effect of Pb^2+^-exposure on adult (PN120) females, with a decrease in one brain region in adult Pb^2+^-exposed males (Figure 5). The most significant changes in brain MOR levels as a result of chronic developmental Pb^2+^ exposure was during the pre-adolescence (PN14) to early-adolescence (PN28) period in which several brain regions exhibited significantly higher levels of MOR in both sexes. The relevance of the increase in brain MOR levels during the pre- and early-adolescence period is of special importance given that this is a critical period where risk-taking behaviors and propensity to addictions is highly likely to occur in humans (Jordan and Andersen, 2017). It has been shown that individuals that engage in drug use during adolescence are more prone to relapse later in life (Jordan and Andersen, 2017) and have increased susceptibility to mental disorders, such as mood disorders (Lutz and Kieffer, 2013). Animal studies have also shown that adolescence exposure to drugs of abuse impacts behavioral, affective, and cognitive functions long after the period of adolescence (Spear, 2016).

Opioid receptors are distributed throughout the nervous system and are responsible for responses to pain stimuli and stress and influence reward processing and mood (Gerrits et al., 2003) and are prominently expressed in areas of the brain associated with addiction circuits (Nestler, 2004). The opioid system comprises three G-protein coupled receptors known as mu- delta- and kappa-opioid receptors and they are stimulated by their respective endogenous opioid peptides (Kieffer, 1995). In particular, MOR is responsible for the analgesic, rewarding, and dependence effects of morphine (Matthes et al., 1996). The MOR also mediates the biological effects of other drugs that are used clinically and have abuse liability such as oxycodone, methadone, heroin, and fentanyl (Darcq and Kieffer, 2018). Therefore, the changes that we have described in the present study provide evidence that Pb^2+^, an environmental pollutant that is ubiquitous in the environment, is able to alter MOR levels and sensitize addiction circuits with the potential for predisposing subjects to substance abuse and risk behaviors. The latter is consistent with human studies indicating that childhood Pb^2+^ intoxication is associated with a proclivity to drug addiction, antisocial behavior, and juvenile delinquency (Carpenter and Nevin, 2010; Fishbein et al., 2008; Needleman et al., 1996; Wright et al., 2008).

While the present study is the first of its kind to examine the effect of chronic developmental Pb^2+^ exposure using a life course approach in male and female rats, it has the limitation that we did not examine the levels of endogenous endorphins (endogenous ligand) or the behavioral correlates associated with increased brain MOR levels in Pb^2+^-exposed animals. Previous studies have shown that a similar chronic Pb^2+^ exposure paradigm (did not provide blood Pb^2+^ levels) was associated with decreased β-endorphin and increased [^3^H]-naloxone binding in the whole of brain at embryonic day 16 (Baraldi et al., 1988). In the same study, they also examined the levels of β-endorphin (hypothalamus) and [^3^H]-naloxone binding (brainstem and hippocampus) at PN7, PN14, PN40, and PN 60. They found a similar effect as in whole brain from embryonic 16 animals, that is, decreased in β-endorphin in the hypothalamus and increased [^3^H]-naloxone binding in the brainstem but not in the hippocampus. Importantly, they found that at PN40, Pb^2+^-exposed rats had decreased sensitivity to pain stimuli (increased time in hot plate test) when compared to control animals and this effect was more dramatic after morphine administration (Baraldi et al., 1988). The effect of morphine in Pb^2+^-exposed rats to pain persisted long after cessation of Pb^2+^ exposure. Other studies using a perinatal Pb^2+^ exposure paradigm provide further evidence of disruption of the opioid system and suggest a possible link between Pb^2+^ exposure and opioid addiction (Kitchen and Kelly, 1993; Rocha et al., 2004; Valles et al., 2003).

## 5. Conclusion

In summary, we show that chronic developmental exposure to moderate levels of Pb^2+^ is able to disrupt the ontogeny of MOR in the male and female rat brain with important implications to opioid use and abuse. Future studies will examine the impact on behaviors mediated via Pb^2+^-influenced changes in brain MOR levels.

## CRediT authorship contribution statement

Damaris Albores-Garcia: Methodology, resources, validation, investigation, formal analysis, writing-original draft and visualization.

Jennifer L. McGlothan: Methodology, investigation and resources. Zoran Bursac: Formal analysis.

Tomas R. Guilarte: Conceptualization, methodology, writing-review and editing, supervision, project administration and funding acquisition.

## Acknowledgements

we thank Deborah Brooks for assistance in the husbandry of animals and Ana G. Sanchez.

